# The influence of scaffold deformation and fluid mechanical stimuli on bone tissue differentiation

**DOI:** 10.1101/2024.02.29.582717

**Authors:** Laia Moliner, Carlos Ruiz Wills, Oscar Camara, Andy L. Olivares

## Abstract

Bone tissue engineering enables the self-healing of bone fractures avoiding the graft surgery risks. Scaffolds are designed to transfer global mechanical load to cells, and the structure-blood flow interaction is crucial for cell differentiation, proliferation, and migration. Numerical models often consider the effect of solid on the fluid or vice-versa, nevertheless, fluid-structure interactions (FSI) are not usually explored. The present study aims to develop in-silico FSI models to evaluate tissue differentiation capability of different scaffold designs. This is accomplished by analyzing the relation between scaffold strain deformation and fluid mechanical stimuli developed at the cell microscopic level. Cubic regular structures with cylinder and sphere pore based of 60%, 70% and 80% porosity were modelled in finite element analysis. Static or dynamic compression and inlet steady state or transient state fluid profile were considered. Fluid-structure interactions have been performed, and cell differentiation studies considering the octahedral shear strain and fluid shear stress have been compared. Results indicate that high porous scaffold with low compression and fluid perfusion rates promote bone tissue proliferation. Moreover, mechanical stimulation seems to help bone formation and to inhibit cartilage phenotype. Results showed that neglecting the interaction between the scaffold and fluid flow could lead to substantial overestimation of bone differentiation. This study enhances our understanding of the role of dynamic mechanical simulations in tissue formation; allowing the improvement of scaffold design to face complex bone fractures.

## 1 Introduction

The bone can repair itself when a fracture occurs. The healing process restore bone original strength and structure. However, when bone defect area is large, e.g. 2 cm or more, the healing mechanism might not be sufficient to regenerate the bone tissue effectively. [22]. Traditionally, clinicians use bone transplantation as a treatment. Insufficient bone source, immune rejection, secondary surgery, or cross-infection are scenarios that occurs in bone transplantation [14]. Lately, bone tissue engineering (BTE) has risen as an alternative technique to deal with large bone defects.

BTE creates new bone tissue in a laboratory setting to repair or replacement damaged or diseased bones [4]. The process involves porous structures known as scafolds that serves as temporally replacement for tissue extracellular matrix (ECM) for mesenchymal stem cells (MSCs) differentiation, proliferation, and migration [1, 22]. Persson et al. (2006) showed the capability of a three-dimensional scaffold to promote the differentiation of MSCs into osteoblast lineage and bone mineralization [20]. Poly Lactic-co-Glycolic Acid (PLGA) and collagen scaffolds have also obtained successful results on in vitro differentiation in bone tissue [23]. However, artificial ECM should be a balance of mechanical function with biofactor delivery, providing a sequential transition in which regenerated tissue assumes function as the scaffold degrades [2, 11].

BTE faces some challege for its regular clinical application. One of the major problems is stem cell differentiation into osteogenic lineage during and after scaffold implantation surgery [4]. Cell differentiation is directly related to the mechanical stimuli received by the cell through the scaffold. Recent studies showed that MSCs differentiation towards osteogenic lineage is regulated by external mechanical stimulation [4]. These external inputs can be caused by microdeformation of the perilacunar bone matrix and by shear stresses generated due to canaliculi fluid flow [4]. Therefore, it is essential to understand how a scaffold transmits fluid flow stresses and mechanical deformations to its surfaces. Experimental settings might be complex, expensive and it is difficult to evaluate specific local data. As such, computational approaches have risen in the recent years [24].

Computational Fluid Dynamics (CFD) have been applied to link scaffold configuration to fluid shear stress (FSS) behavior in BTE [3, 17, 19]. Several studies have been conducted to correlate scaffold characteristics with FSS outcomes. Ali et al. [3] used CFD analysis to study the effect of surface roughness on scaffold permeability and fluid flow-induced FSS and concluded that high roughness decreased FSS. Ouyang et al. [19] evaluated the effect of pore size on hydro-mechanical properties and stated that flow velocity linearly increased with the pore size. Recently, Fluid-Structure Interaction (FSI) models have be introduced to address the impact of mechanical compression on FSS calculation [27]. Zhao et al. [27] used CFD and FSI models to quantify the FSS of different scaffold geometries (architecture, pore size, and porosity) under the combination of fluid perfusion and compression loading scenarios. They showed that the combination of both stimuli might cause the amplificatrion of the FSS. Fu et al. [9], also a combination of CFD and FSI approaches to evaluate the permeability and FSS ranges of different scaffold configurations applying steady state mechanical inputs. Their result indicate that the pore size of the scaffold increases, its permeability increases and the FSS decreases [9]. However, these studies only consider the FSS results to evaluate scaffold viability, without also considering the direct impact that scaffold shear strain might have on cell differentiation.

The main objective of the present study is evaluate both transient mechanical and fluid stimuli in the ideal seeding scaffold for different scaffold design. Hence, the primary purpose is to perform a fluid-structure analysis to evaluate their potential for bone formation. The study was focuses on the optimal configuration promote a cell differentiation in order to restore a bone lesion.

## 2 Materials and Methods

### 2.1 Scaffold design

Two cubic regular architectures were selected following the design proposed by Fu et al. [9]: cylinder pore-based (C), and sphere pore-based (S). The design size of each model was 2 × 2 × 2 mm, which was generated by coupling 0.5mm basic units (Fig. 1 and Fig. A1) using the software Fusion360 (Autodesk, Inc). Three levels of porosity were considered: 60, 70, and 80 % based on previous experimental studies [26]. In total, 9 different geometries were evaluated with disparate characteristics (Table 1).

**Figure 1.**
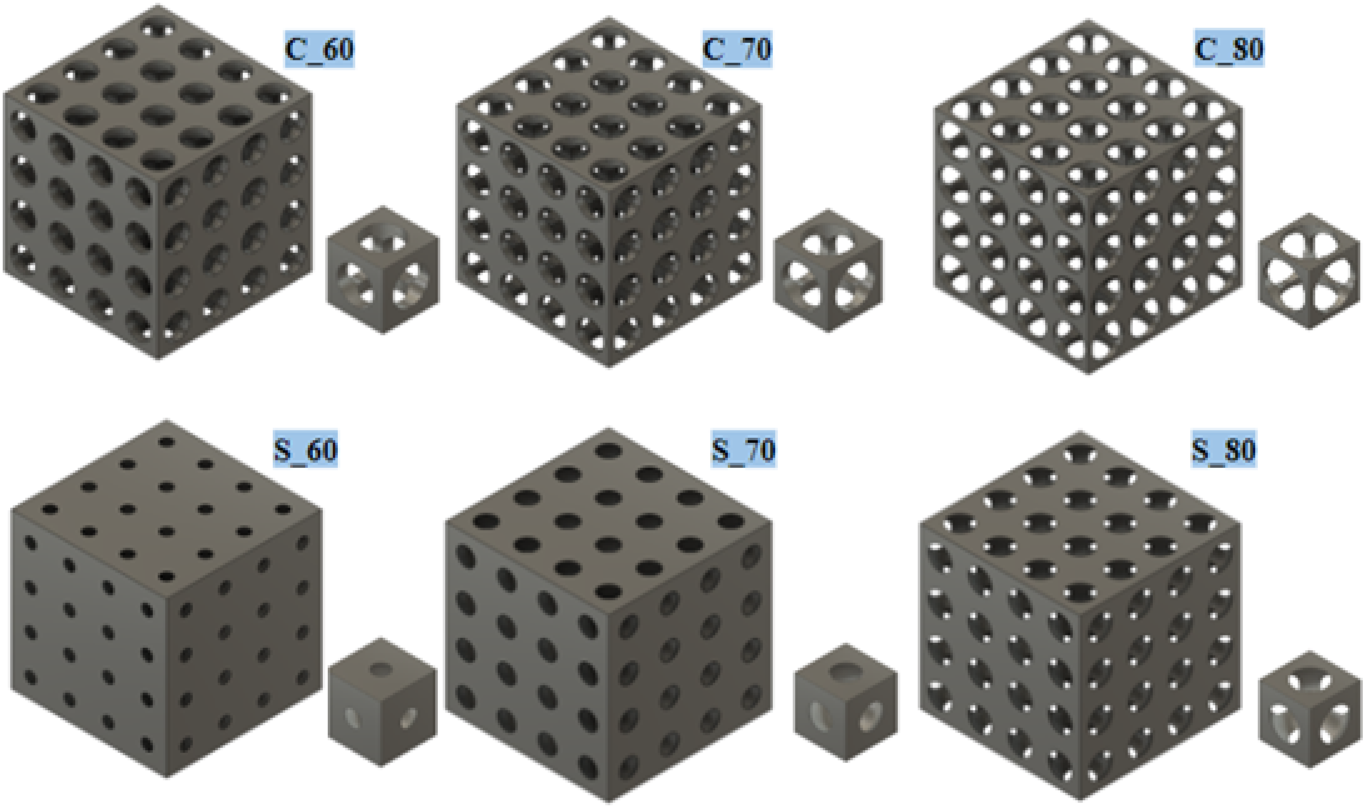
Computer-aided design (CAD) diagrams of the scaffolds. Left first row: Basic cylinder porous units with 80, 70, and 60% of porosity. Left second row: Basic sphere porous units with 80, 70, and 60% of porosity. Middle: 60% cylinder porous scaffold. Right: 60% sphere porous scaffold.

**Table 1.** Properties of the six scaffolds denoted as the type of porous followed by low bar and degree of porosity. For example, the cylinder porous scaffold with a porosity of 60% is named C 60.

| Scaffold model | Porosity (%) | Porous diameter (mm) | Surface area (mm<sup>2</sup>) |
| --- | --- | --- | --- |
| C_60 | 60 | 0.32 | 57.18 |
| C_70 | 70 | 0.36 | 52.42 |
| C_80 | 80 | 0.41 | 43.69 |
| S_60 | 60 | 0.14 | 70.56 |
| S_70 | 70 | 0.23 | 64.27 |
| S_80 | 80 | 0.29 | 57.13 |

### 2.2 Three-dimensional meshing

The scaffolds generated in section 2.1 were discretised in Abaqus 2018 (Dassault Systèmes) to create an optimal 3D tetrahedral mesh for the solid part. The mesh size was established at 0.05 mm considering the computational cost, the possible approximation error, and knowing that cells diameter oscillates around 0.01 mm [8]. A fluid boundary was created with the insertion of an external cube of 2.1 mm of dimension using Meshmixer (Autodesk, Inc). The external cube and the scaffold were aligned.The space between solid and external fluid boundary was fulfil with 3D tetrahedral elements via Gmsh [10]. Finally, the final mesh quality was evaluated in Abaqus 2018 (Dassault Systèmes).

### 2.3 Structural model

Computational Solid Mechanics (CSM) simulations were performed for each scaffold using Abaqus 6.12 (Dassault Systèmes). Poly-D, L-lactic acid (PDLLA) was selected for all scaffolds. PDLLA was considered as linear elastic, with a density of 1260 Kg*/*m^3^ [15], a Young’s Modulus of 2 GPa [25], and a Poisson’s ratio equal to 0.331 [6].

CSM simulations consisted in the applications of a compressive load at the upper surface of the scaffold with a rate of 0.016 mm/s (Fig. 2). The lower surface of the scaffold was fixed in all directions. The maximum compression achieved in the simulations was equal to 5 % displacement relative to the height of the scaffold [13],i.e. a total displacement of 0.1 mm. The models were evaluated with two different displacement profiles: a) static-state compression, and b) dynamic-state compression. The static-state simulations considered a constant displacement of 0.1 mm by 0.1s. In a dynamic-state simulations, a load-unload cycle for 0.2s was performed, i.e. a compression until 0.1 mm in the first 0.1s, and removing the 0.1 mm of compression in the following 0.1s.

**Figure 2.**
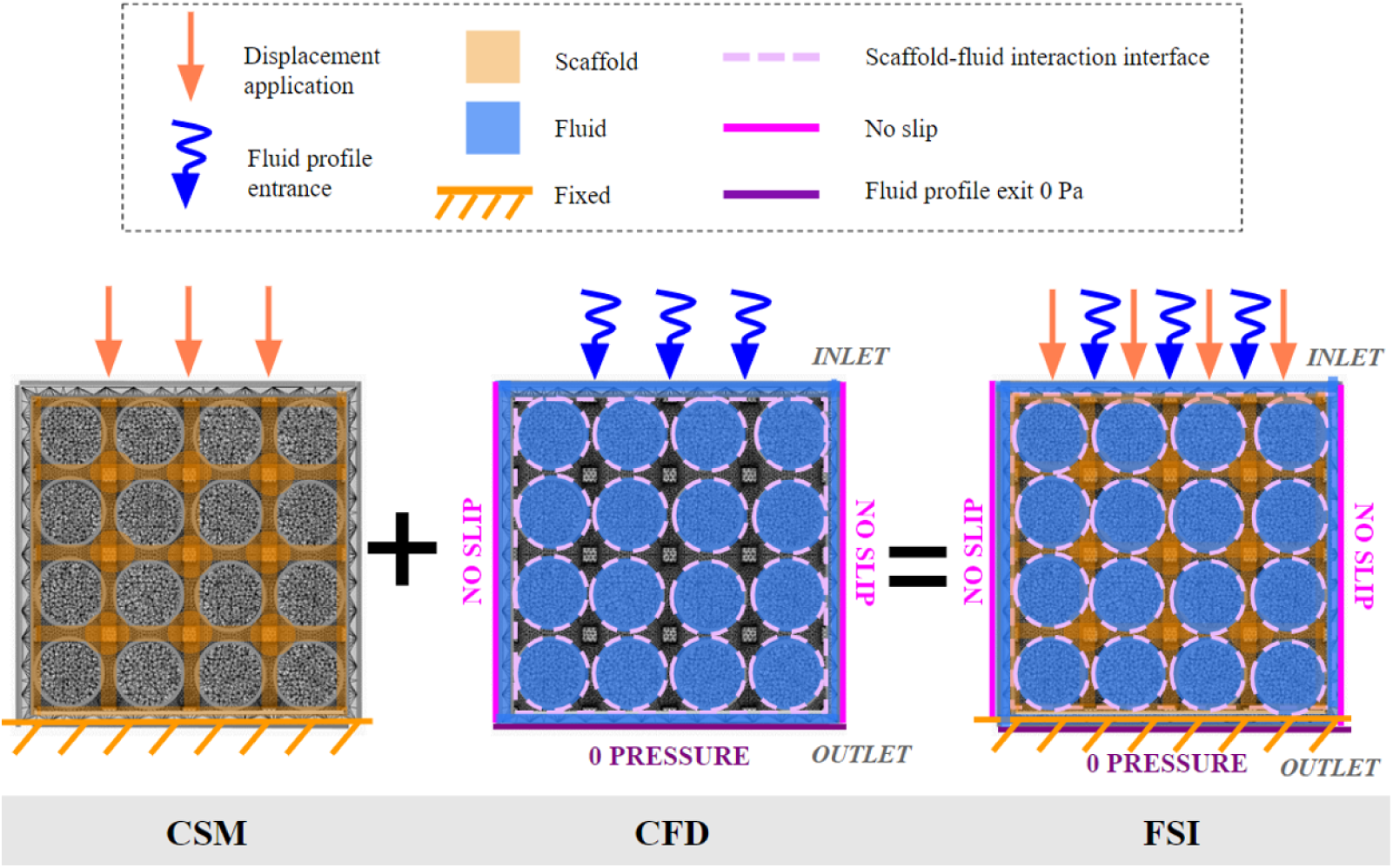
Scheme exhibiting the boundary conditions applied during the simulations performed during the study. Left model: For Computational Solid Mechanics (CSM) simulations, a displacement was applied on the top part of the scaffold (coral arrows) and the inferior part was fixed (orange marks). Middle model: In the case of Computational Fluid Dynamics (CFD) simulations, a fluid profile was applied on the inlet (blue arrows) and a zero pressure was established in the outlet (dark purple). Moreover, no-slip conditions were considered for the left and right exterior nodes (pink line), and the interaction interface between the scaffold can be observed in light purple dotted lines. Right model: For the Fluid-Structure Interaction (FSI) simulations all the boundary conditions mentioned for the CSM and CFD were applied simultaneously.

The shear strain components and the reaction force were extracted from the simulation. The Effective Young’s modulus (*E_f_* ) [17] was calculated using Equation 1 and the shear strain (SS) was computed using Equation 2 [13, 17, 21].

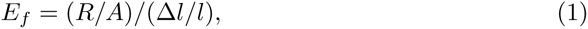

where R is the reaction force, A is the cross-section area of the scaffold, and Δ*l/l* is the axial strain [17].

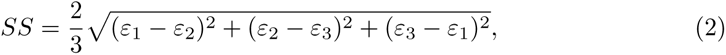

were *ε*_1_, *ε*_2_ and *ε*_3_ are the principal strains (maximum, medium and minimum, respectively) [17].

### 2.4 Fluid model

Computational fluid dynamics (CFD) simulations followed the Navier-Stokes equations. The fluid selected was Dulbecco’s modified Eagle medium (DMEM). The DMEM was modeled as a Newtonian fluid with the transient state viscosity of 1.45 *×* 10*−*3 Pa s, a density of 1000 Kg*/*m^3^and assuming a laminar flow [17].

An inlet fluid velocity profile was homogenously applied for 1 mm/s in the inlet and a zero pressure assumed at the outlet [17, 19](Fig. 2). A non-slip wall boundary condition was applied to the exterior fluid walls and the interior fluid walls in contact with the scaffold. Moreover, two fluid velocity profiles were tested to examine the different responses to the stimulus. On the one hand, a steady-state fluid simulation with a constant fluid velocity equal to 1 mm/s [17] for 0.1s was applied at the inlet. On the other hand, a transient state simulation was performed using a fluid velocity cycle of 0.2s. The cycle consisted in increasing a fluid velocity of 1 mm/s for 0.1s, and decreasing again until 0 mm/s.

### 2.5 Fluid-Structure interaction model

After simulating the solid and the fluid phases separately, the real-time interactions between the scaffold and the fluid were evaluated through fluid-structure interaction models. Therefore, to be as accurate as possible to reality, the models were based on a specific bioreactor design. Within this bioreactor the fluid could freely transport through the scaffold ends under mechanical displacement, and the scaffold is laterally confined. Furthermore, four cases were studied: a) static fluid profile without compression, b) transient fluid profile without compression, c) transient fluid profile with static compression, and d) transient fluid profile with dynamical compression. The strain components, FSS, fluid velocity, and fluid pressure were computed using Abaqus 6.12 (Dassault Systèmes). The SS was also calculated using equation 2.

As previously mentioned, the models follow a particular bioreactor design, hence the boundary conditions were established as can be seen in Figure 2. For the fluid model: the outlet was defined a zero pressure; the particular fluid profile was homogenously applied along the Y-axis; and a non-slip condition was applied to the exterior walls. For the solid model: the inferior part was fixed, and the superior part was displaced uniformly along the Y-axis with a steady or dynamic state compression profile. Moreover, both models were connected for the interaction surface, defined as the interior fluid wall and the exterior scaffold wall. However, the top and bottom surfaces have to be excluded from the interaction surface since there were certain restrictions and displacement already implemented.

### 2.6 Mechanobiology stimuli

The cell diffentiation evaluation at the scaffold surface followed the method proposed by Olivares et al. 2009 [17]. In brief, the scaffold surface percentage associated with bone, cartilage, or fibrous tissue is described in terms of inlet parameters. The mechano-regulation theory proposed by Prendergast et al. 2002 [12] was modified to include the SS from the solid phase and the FSS from the liquid phase. As a result, the SS and FSS obtained from the simulations were used to compute the stimuli S on the scaffold wall by using Equation 3. If 0.01 *≥ S*, the stimuli are too low to induce any type of tissue phenotype, and if *S ≥* 6 the stimuli will be too high to induce differentiation. If 3 *≥ S ≥* 1, then cartilage tissue differentiation occurs, 6 *≥ S ≥* 3, then fibrous tissue differentiation occurs, and if 1 *≥ S ≥* 10*^−^*^2^ bone differentiation occurs [12, 14, 17]

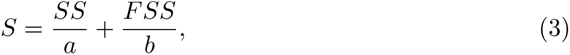

Stimuli S computed using the octahedral Shear Strain (SS) and the Fluid Shear Stress (FSS). Where a is equal to 0.0375% and b is equal to 10mPa [17].

Different cell differentiation analyses were executed (Fig. 3). Firstly, the SS obtained from the CSM model (for the static and dynamic cases) was evaluated to extract the element ratio differentiation, only considering the x-axis classification. The same was done for the FSS resulting from the CFD model (for both fluid profiles) but only considering the Y-axis. Secondly, to acquire the cell differentiation distribution for both parameters, a node coupling was performed to determine the common nodes, and then the stimuli S were calculated for each node. Thirdly, the FSS obtained from FSI models that did not include displacements was analysed only by contemplating the Y-axis. Subsequently, these results were compared to the CFD results but only considering the shared nodes (as the interaction surface of the FSI did not involve the inferior surface of the scaffold). Finally, for the FSI simulations with deformation, a coupling of FSS and SS results was performed.

**Figure 3.**
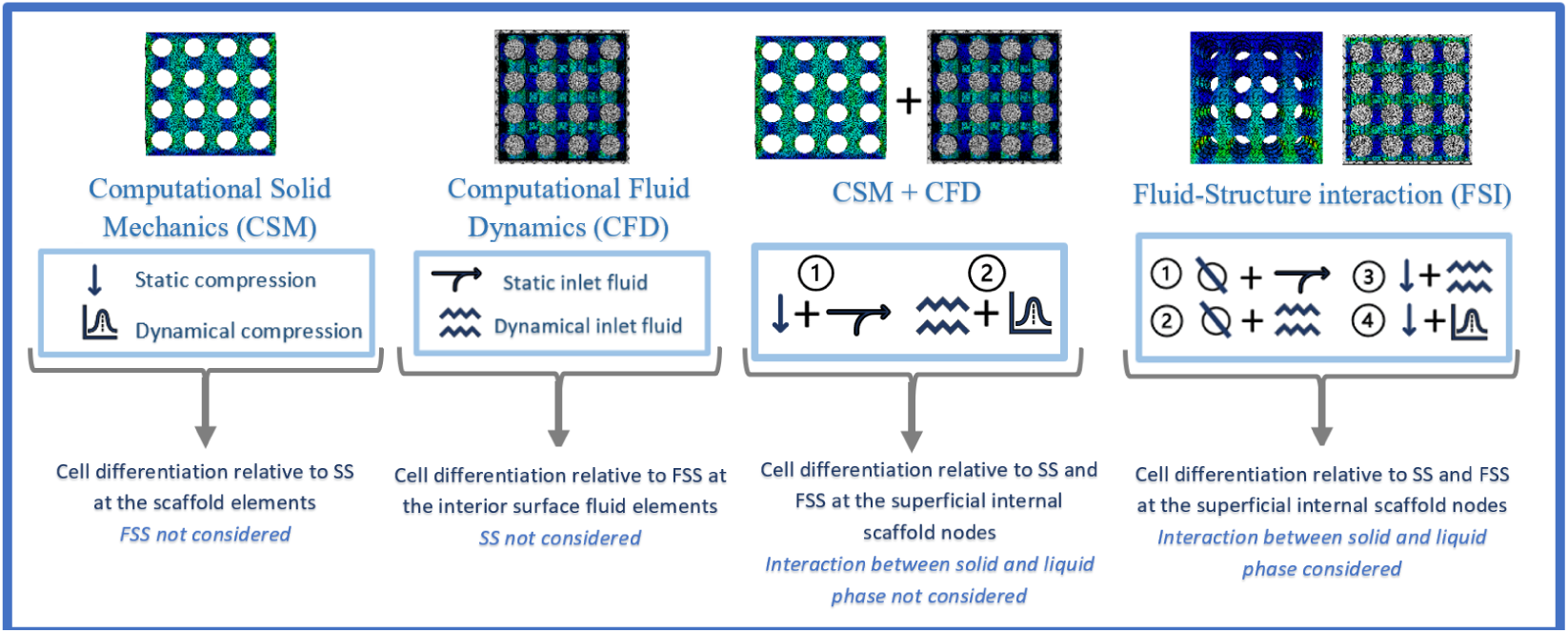
Concept mapping explaining the four computational approaches used in the study to determine the bone differentiation of the different scaffold conformations. (1) CSM where SS results were contemplated but the fluid phase was excluded. (2) CFD where FSS results were considered but the solid phase stimulation was excluded. (3) The correlation between the CSM and CFD results was performed with the solid and the fluid phase but with no interaction between them. (4) FSI model in which the solid and fluid phases were taken into account and there was an interaction between the surfaces.

## 3 Results

### 3.1 Mechanical properties of the scaffolds

Table A1 shows that for higher porosities the *E_f_* decreased. The maximum value of *E_f_* was obtained for the configuration C 60 and the minimum for S 80. Moreover, the cylinder pore-based scaffolds presented higher *E_f_* values than the scaffolds sphere pore-based.

The percentage of elements under compression is notably higher than the elements under tension, and the differences between porosities are minimal (Fig. 4). At 60% of porosity, the maximum elements under compression were achieved for both scaffold types. In fact, for the C 60 around 87.8% elements were under compression and for S 60 around 89.3% (Fig. 4). The lower values were achieved at 80% porosity with 77.2% and 84% of elements under compression for C 80 and S 80, respectively (Fig 4). Furthermore, at the end of CSM, the stress distribution (compression and tension) were localized in horizontal areas between the pores for both cilindrical and spherical scaffolds (Fig. A2).

**Figure 4.**
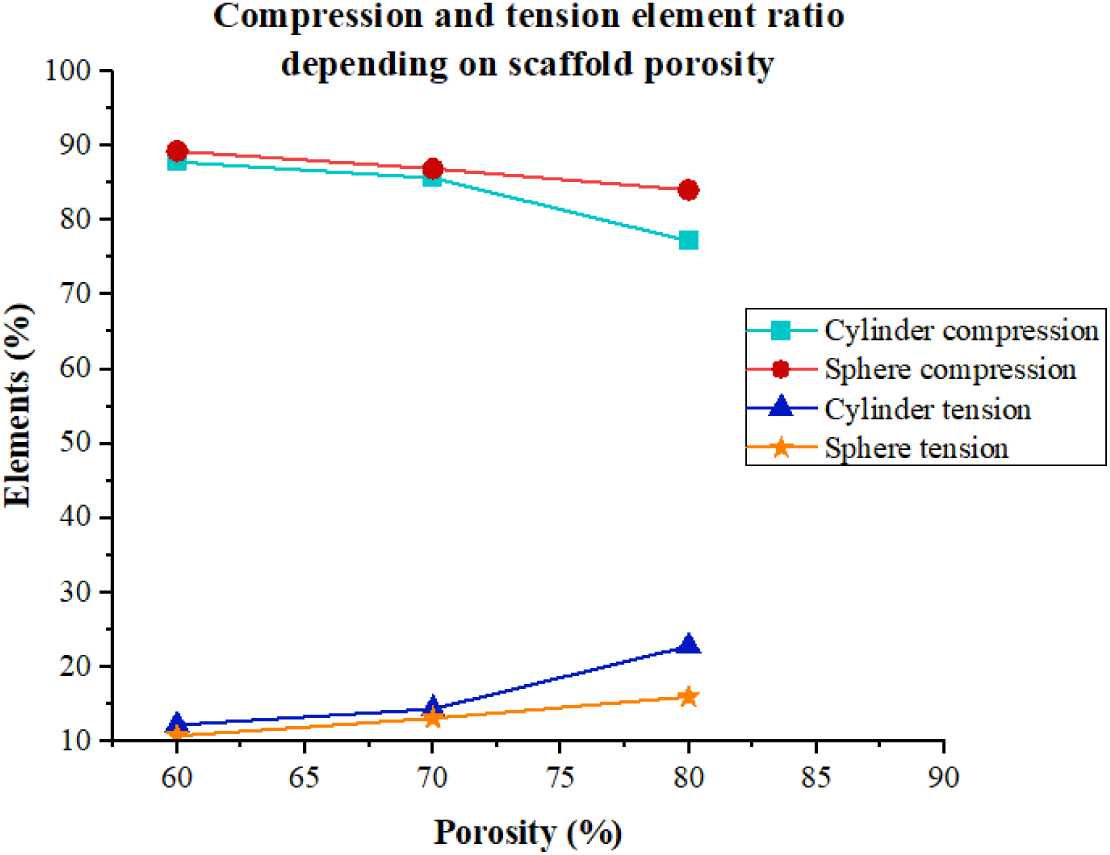
Diagram showing the ratio of elements (y-axis) for the cylinder model under compression (light blue) and tension (dark blue), and sphere models under compression (red) and tension (orange) for 60%, 70%, and 80% (x-axis).

### 3.2 Structural model

Bone differentiation was relatively high in static-state simulation for both scaffold configurations. C 80 showed the maximum value with 76% of bone and S 60 the lowest value with 51.4% of bone. Cylinder-shaped pore scaffolds presented higher bone differentiation than spherical-shaped pore scaffolds. In addition, low porosity’s presented less bone phenotype than high porosity’s.

Similar cell differentiation outcomes were obtained with dynamic compression (Fig. 5).The variation in differentiation percentage between cartilage and bone was larger than with static-compression. For low deformation, there was high differentiation of bone, and low differentiation of cartilage and fibrous tissue. However, when the compression stimuli increased, bone tissue decreased and cartilage and fibrous tissue

**Figure 5.**
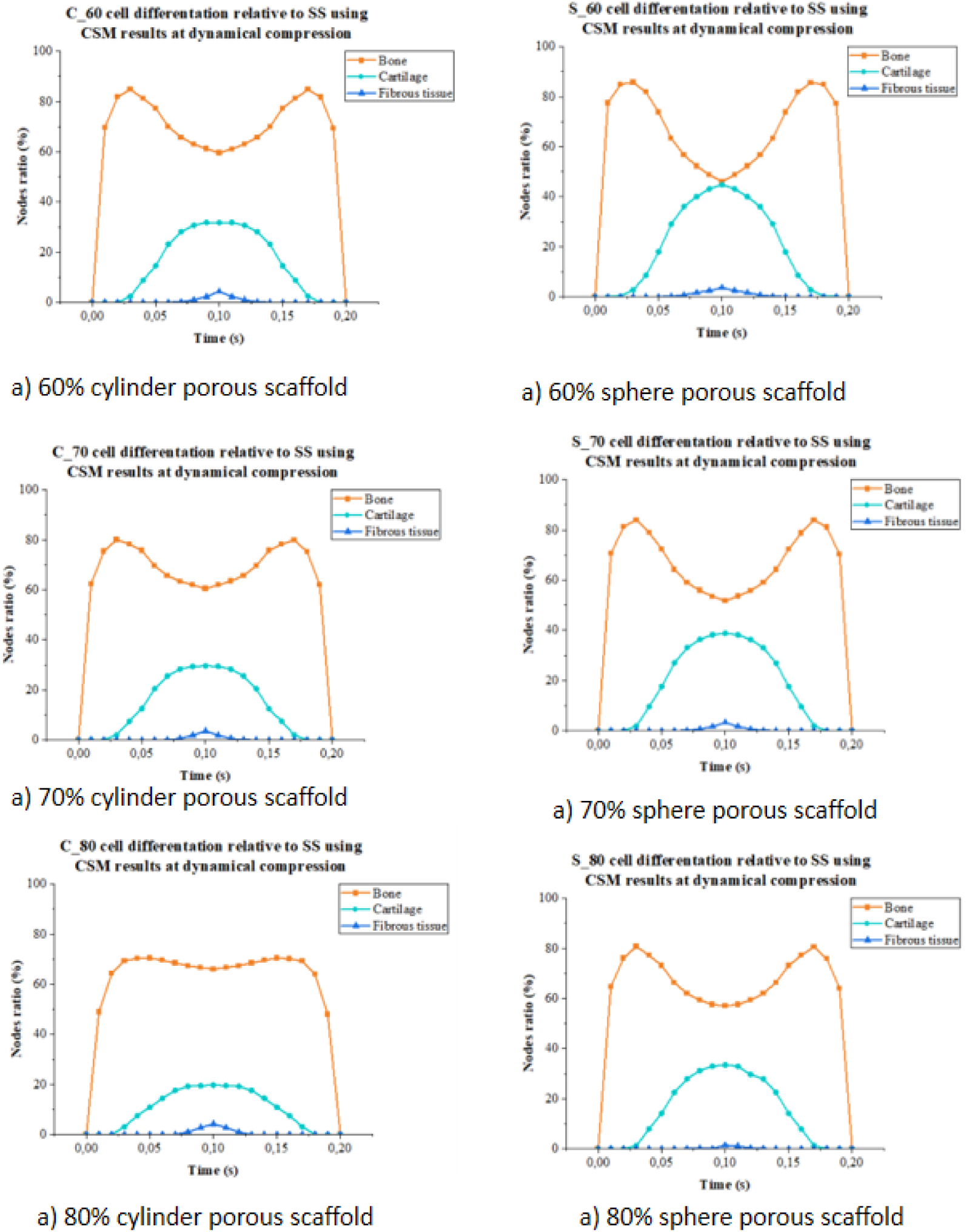
Cell differentiation analysis relative to Shear Strain (SS) results in the superficial nodes using Computational Solid Mechanics (CSM) at dynamic compression, being bone differentiation orange, cartilage differentiation light blue, and fibrous tissue differentiation dark blue.

### 3.3 Fluid model

Low values of bone tissue differentiation were obtained for static fluid profile (Fig. A5). Both scaffold type led to the same outcome with a slightly higher bone differentiation for low porosity. In contrast, the percentage of fibrous tissue differentiation is remarkably high in comparison with bone. Transient state simulations showed a peak of bone differentiation for slow velocity, followed by a peak of cartilage differentiation and a wider peak of fibrous tissue differentiation (Fig. A4). The maximum peak for bone differentiation were 52.45% for C 80 and 44.57% for S 80.

### 3.4 Fluid-Structure model with no interaction

Negligible bone differentiation was obtained for both scaffold types (Fig. A7). However, for low fluid and mechanical stimuli, a small peak of bone differentiation appeared for S 80 and C 80. Yet, most of the stimuli applied were too high to promote tissue formation (Fig. A8).

### 3.5 Fluid-Structure interaction model

Figure A9 shows the cell differentiation of FSI models for transient state fluid profile and no compression. The results showed low bone tissue differentiation for a maximum velocity of 1 mm/s. Figure A6 showed that FSI bone differentiation for slow velocity was slightly lower than CFD differentiation. For the peak a high velocity, differentiation gap between FSI-CFD increased. Indeed, for C 80 the second peak was almost nil, and S 80 showed the highest second peak but lower than CFD. FSI showed that the deformation produced by the fluid on the scaffold increased from top to bottom (Fig. A10).

The results obtained from FSI models steady state for both compression and fluid flow showed no differentiation of any type of tissue for the S 80 scaffold (Fig. A11a). Similar outcomes were obtained with steady compression and transient state fluid (Fig. A11b). Indeed, the S stimuli was too high to promote any tissue differentiation as obtained in section 3.4. SS was concentrated around the scaffold porous when both fluid and mechanical stimuli were considered (Fig. A12).

## 4 Discussion

Fluid-structure analysis on different scaffold configuration showed that cylindrical shape with 80% of porosity is the optimal configuration that might lead to bone cell differentiation enough to repair a fracture. Using both mechanical and fluid stimuli together led to a better representation of the stimuli that occurs physiologically. For example, in Olivares et al. [17], they used the same mechanoregulation theory but separate mechanical and fluid stimuli. Their results suggested that the cell differentiation is more sensitive to the fluid shear stress than to the axial strain. The calculation of both stimuli at the same time, and to allow the communication between them is one of the main advances recommended in this paper. A coupled role of the scaffold matrix and interstitial fluid can increase their osteoinductivity. CSM simulations indicated that all the scaffold geometries might have enough mechanical properties to be considered as a possible bone substitute. As shown in Table A1, all the *E_f_* values obtained are similar to Young’s modulus established for the scaffold material. In Figure 4, most elements in all the scaffolds were under compression, which is positive as it is generally accepted that bone strength is greater in compression than in tension [16].

The cell differentiation study using CSM, confirm that a compression between a range of 0 to 5% is crucial to obtain high percentages of bone phenotype. However, the results also indicate that maximum compression were less favorable for bone tissue differentiation (see Fig. A4). For instance, the S 60 scaffold presented a similar differentiation percentage for bone and cartilage at high deformations (51.4% and 44.8%, respectively), increasing the risk of mechanical failure. It is essential to highlight that cartilage and fibrous tissue do not possess similar mechanical properties as bone; hence the presence of significant regions of these tissues can cause instability and even a second fracture. Therefore, cylinder pore-based scaffolds with high porosity could be more optimal than sphere-based for bone differentiation. Since cylinder scaffolds achieved lower cartilage and fibrous tissue differentiation, the bone values remained more stable in comparison with the sphere scaffolds during the dynamic state deformation. This outcome suggest that cylinder scaffolds are transmitting lower SS stimuli than sphere pore based but high enough to remain within the range needed for bone tissue differentiation. Consequently, since stimulus cannot be controlled in a natural environment, contrary to a bioreactor, it could be more positive to have a scaffold presenting lower and more stable peaks of bone differentiation and lower phenotype presence of other tissue types.

Cell differentiation results obtained for CFD method showed that the bone differentiation is achieved only under low velocities level. Indeed, at very low velocities, a peak of bone phenotype was found, followed by a peak of cartilage phenotype. In addition, an increment of the velocity entailed a notable presence of fibrous tissue differentiation, while bone and cartilage were drastically decreased. CFD results suggest that 1 mm/s is too high to obtain satisfactory rates of bone tissue differentiation. Even so, if only the fluid stimulation is considered, it seems complicated to obtain enough bone differentiation and avoid regions with another tissue type. Since at low velocities, the cartilage tissue is remarkably present using these scaffold designs even at high porosity conformations that seem to obtain the best results.

Furthermore, velocity and bone phenotype can be correlated (see Fig. A6), as high bone differentiation peaks are achieved when the velocity rate is low. A clear example of this is the comparison between the high porous scaffold and low ones; the first type presents lower velocities than the second one (Fig. A6), and accordingly, the ranges of bone differentiation are higher for high porous scaffolds than for low porous scaffolds. However, it is essential to clarify that scaffolds with lower porosities often achieved better bone phenotype results than highly porous ones. However, since these fluid perfusion rates have been confirmed too high to obtain sufficient bone differentiation (percentages are too low), this does not affect the conclusion. Moreover, this clarification agrees with the results of Ouyang et al [19]. and Fu et al. [14], that reported ans increase in FSS when the pore size increases. For this reason, when the fluid perfusion velocity achieves high rates, the FSS values will increase more in high porous scaffolds than in lower ones. Therefore, as Olivares et al. [17] study indicated, high FSS values do not promote bone tissue differentiation.

Nevertheless, a coupling between the mechanical and fluid results was made to obtain a more realistic analysis of the stimulation that mesenchymal stem cells could undergo. The results shown in Figure A7 reiterate the conclusions extracted from the previous models. Only some bone cell differentiation is present when the fluid flow is minimum, and the compression is low due to an excessive high stimulation that inhibits cell differentiation (see Fig. A8). Additionally, the SS analysis reaffirmed that high porous scaffolds present more bone differentiation than low porous. Considering that high porosity scaffolds achieved better results in mechanical stimulation than with fluid simulation, it is possible to conclude that compression deformation might be fundamental to obtain bone differentiation at low velocities and to inhibit cartilage differentiation. Cylinder pore-base scaffolds achieves more bone differentiation than sphere pore-based scaffolds. Yet, neither of them with the velocities considered in this study could obtain significant results to restore a bone fracture. Despite these results, it could be interesting to analyze these scaffolds with the same deformation range at low velocities to be capable of discerning the actual application of these scaffold designs.

Most studies until now have evaluated the scaffold performance and predicted its adequate application in BTE, contemplating the solid and liquid phases as independent parts [14, 27]. Accordingly, even though they present a FSI study, there is no real interaction between the two parts within the simulation. What is commonly done is to perform a CFD simulation, assuming that due to the high Young Modulus scaffold, there cannot exist any deformation of the scaffold caused by the fluid [27]. Then, the pressure results obtained from the CFD are introduced to a CSM simulation, and the compression is done. Knowing this strategy, a critical objective of the study was to verify if considering the interaction scaffold-fluid, the results remain the same or not. Nonetheless, as it is shown in Figure 6, it is clear that the results obtained were quite different.

**Figure 6.**
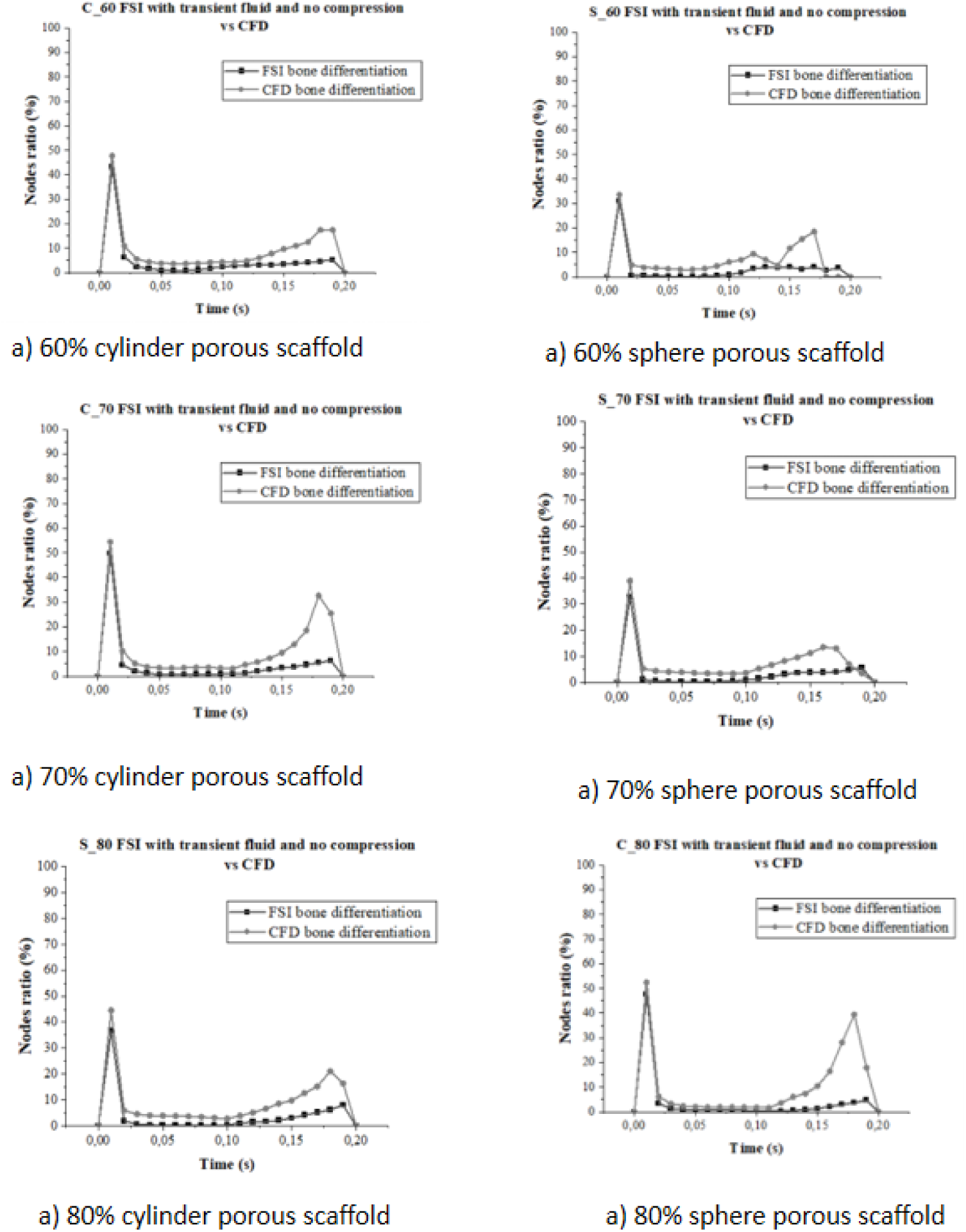
Bone differentiation comparison analysis between Fluid Shear Stress (FSS) results obtained using Fluid-Structure Interaction (FSI) model with transient state fluid and no compression (black), and FSS results obtained using Computational Fluid Dynamics models (grey). increased.

The significant difference between a CFD and FSI model is that the first does not contemplate any interaction between the fluid and the scaffold wall, as the wall has no properties and has a 0 velocity. On the contrary, the FSI model, in this case without compression to make possible the comparison with CFD, allows the interaction between the fluid and the solid, conferring specific properties to the scaffold walls and not applying a zero velocity requirement. Figure 6 shows that CFD and FSI seem to have similar behavior. However, when time increases, the differences increases. This differences can be explained by the fact that the scaffold wall was defined as no-slip in the CFD cases, which means that the wall has zero velocity. Contrarily, the FSI model allowed the deformation of the PDLLA scaffold due to the fluid (see Fig. A10). Indeed, when the fluid impacts the wall, the velocity of the fluid will change, adopting the velocity of the wall during the deformation. These changes are translated into a smaller bone differentiation percentage that will be more affected depending on the scaffold deformation. The results suggest that the studies in which only CFD models are applied might lead to an overestimation of bone tissue differentiation. For this reason, FSI models that allow the interaction of the fluid and the scaffold are necessary for evaluating the scaffold designs for the BTE application.

The scaffold S 80 obtained the best bone differentiation outcomes in the FSI analysis with no compression and transient state fluid. Due to this, it was thought that this design could suffer minor deformation due to the fluid, so it could be interesting to see the results of adding an external compression. However, as mentioned before, the stimulation was too high to promote tissue differentiation. Therefore, it could not be possible to compare the results with those obtained correlating the CSM and CFD results since the differences could not be appreciated.

After performing all the simulations, it can be verified that the horizontal areas between the pores presented more stimuli (Fig. A12). In addition, Figure A3 shows specific fluid flows that collide and impact the wall scaffold. This observation could entail cell seeding in this area, indicating that cells could be attached in these zones where more mechanical and fluid stimulus has been found. This is a decisive observation since there is more certainty that external stimulation will affect mesenchymal cells.

The current study have different limitations. The first are the mechano-regulation theories, biology of bone growth (regeneration), cellular differentiation assumption, amount others. Despite the knowledge of these may help to improve our understanding of bone tissue regeneration, it is complicated numerically replicate all phenomenon take place in vivo or in vitro. As such, numerical models need always simplifications. For cell seeding, it was assumed that the scaffolds were one hundred percent covered with cells, which is why it is overestimated; it is recommended to do a local cell seeding study, such as the one proposed by Olivares et al 2012 [17, 18].

The controlled distribution of the pores might guide the controlled distribution of the differentiated tissue; helping integration in places with certain complexities. However, according to the simulations proposed in this article and in previous ones, it is not possible to know the time it takes to maintain this level and type of stimuli to obtain the desired tissue. We believe that the effectiveness of the application of these techniques will come when we are able to replicate the time in which the stimulus can be withdrawn or changed to another stimulus, once the cells are differentiated and the tissue stabilization process can continue. The quantity of bone tissue differentiated depend not only the scaffold design, but also the bioreactor conditions [7] and to know well the outcome of vitro fabricated scaffolds is possible to recreate proper cell microenvironment during cell differentiation.

FSI models show lower values in bone differentiation. Is important to understand if is really the motion of fluids affects the deformation of structures and vice versa, that is difficult in rigid scaffolds. The material using for the scaffold can be considering rigid in comparison with another biological materials and also is in our proposal a imitation of bone. However, our interest is to know what happens in term of stimuli in the scaffold surface and how simulating the interactions its possible improve or not the results. The coupling is achieved through iterative algorithms, and information is exchanged between the two domains at each time step to ensure a consistent solution, that point is critical and need to be solve in time convergence study. In some previous studies [5, 17], it were sufficient or more efficient to study the fluid and solid structures independently. Nevertheless, our opinion is its needed the validation of process to confirm what is the best way to simulate the differentiation phenomenon. Therefore, the scaffold performance must be evaluated with dynamic mechanical and transient state fluid profile stimulation, and the interaction between the solid and fluid phases must be considered.

A example of one-way FSI model for predicting cell differentiation at the surface of porous hydrogel scaffold, under mechanical stimuli was presented by Azizi et al [5]. This study considers both stimulation: due to mechanical deformation, and due to compression that induced fluid flow during dynamic compressive stimulation. The outcome is very useful for hydrogel application but we consider to mimic better the nature is important to understand the two-ways simulation.

## 5 Conclusions

In this study, FSI model focus on cell differentiation was compared with others two models, shown the differences between each combination. High porous scaffolds with low compression and slow fluid perfusion rates promote bone tissue phenotype. A direct mechanical stimulation has been found to stimulate bone differentiation and inhibit cartilage phenotype. Accordingly, to achieve a correct approximation to elucidate if a scaffold is optimal for successful bone fracture healing, a study mimicking the closest possible to the natural environment is needed.

## Author contribution

Conceptualization, A.L.O. and C. R. W.; methodology, L.M., C.R.W, A.L.O ; software, L.M., C.R.W; formal analysis, L.M., C.R.W, A.L.O; investigation, L.M., C.R.W, A.L.O; resources, C.R.W, A.L.O, O.C. ; writing—original draft preparation, L.M., C.R.W, A.L.O, O.C; supervision, C.R.W, A.L.O . All authors have read and agreed to the published version of the manuscript.

## Acknowledgement

This research received no external funding

### A Appendices

**A.1.**
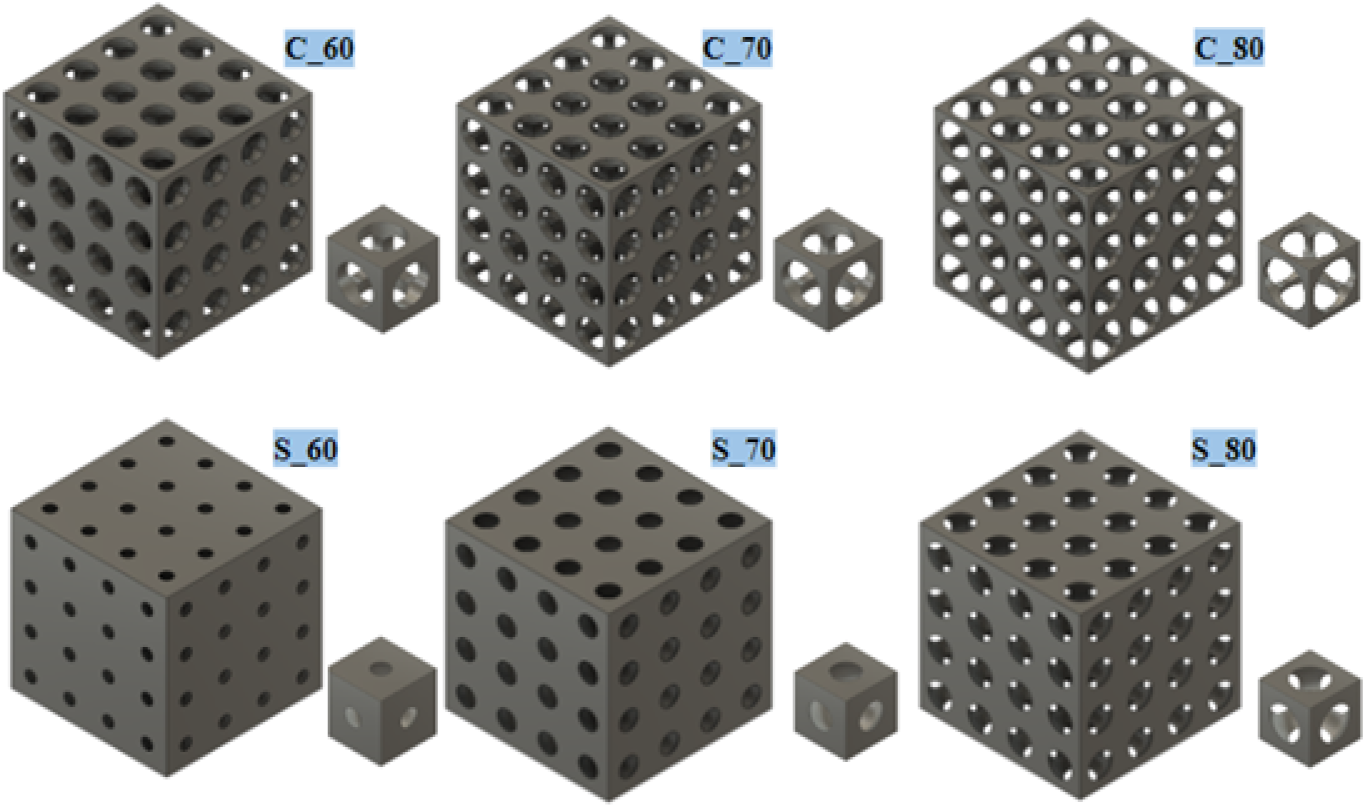
Scaffolds were designed with three different porosities (60, 70 and 80%) and two geometrical pore configurations; cylindrical and spherical.

**Table A.1.** Effective Young’s Modulus of each scaffold design computed with the results obtained from Computational Solid Mechanics at static compression and implementing Equation 1

| Effective Young's Modulus (GPa) |  |  |  |  |  |
| --- | --- | --- | --- | --- | --- |
| C_60 | S_60 | C_70 | S_70 | C_80 | S_80 |
| 0.992 | 0.684 | 0.771 | 0.582 | 0.625 | 0.516 |

**A.2.**
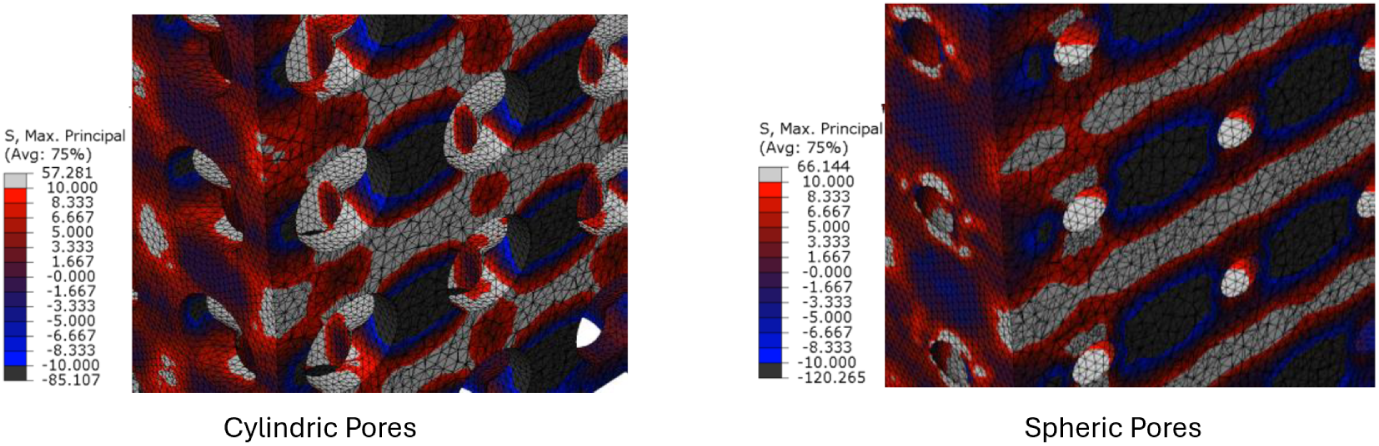
View of the 60% porosity scaffolds showing the absolute stress and strain scale, highlighting the compression areas in blue and black and the tension areas in red.

**A.3.**
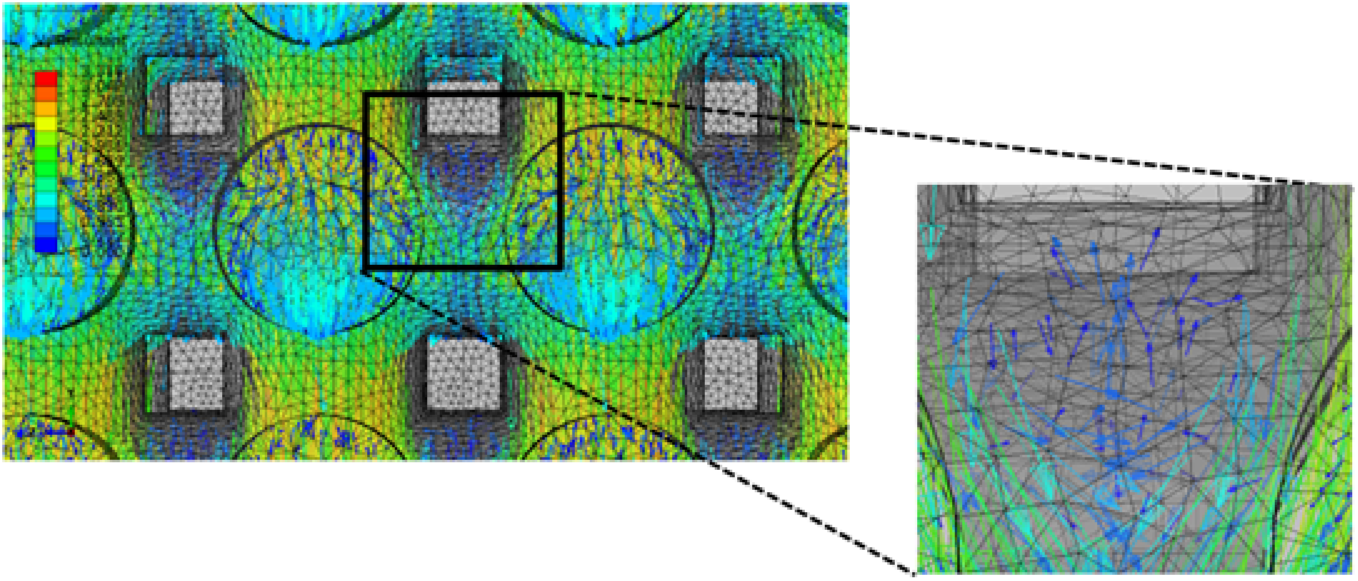
Computational Fluid Dynamics velocity vector color map results at maximum fluid perfusion equal to 1 mm/s. 60% (left) and 70% (rigth) . Furthermore, a zoom is conducted to show how the fluid flows collide and then impact the wall of a 70% cylinder porous scaffold.

**A.4.**
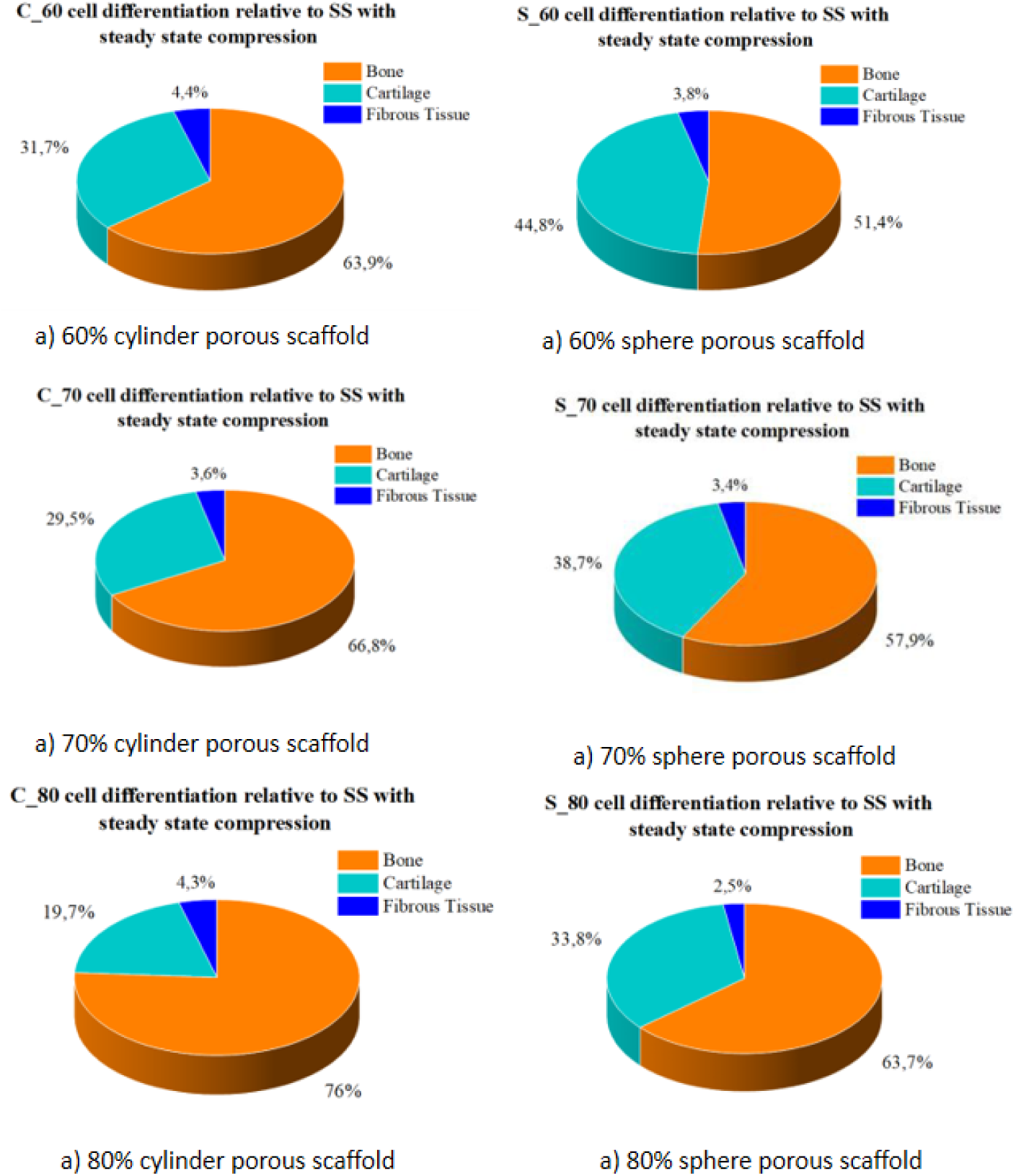
Cell differentiation analysis relative to shear strain (SS) results in the superficial nodes using Computational Solid Mechanics (CSM) at steady state compression, being bone differentiation orange, cartilage differentiation light blue, and fibrous tissue differentiation dark blue.

**A.5.**
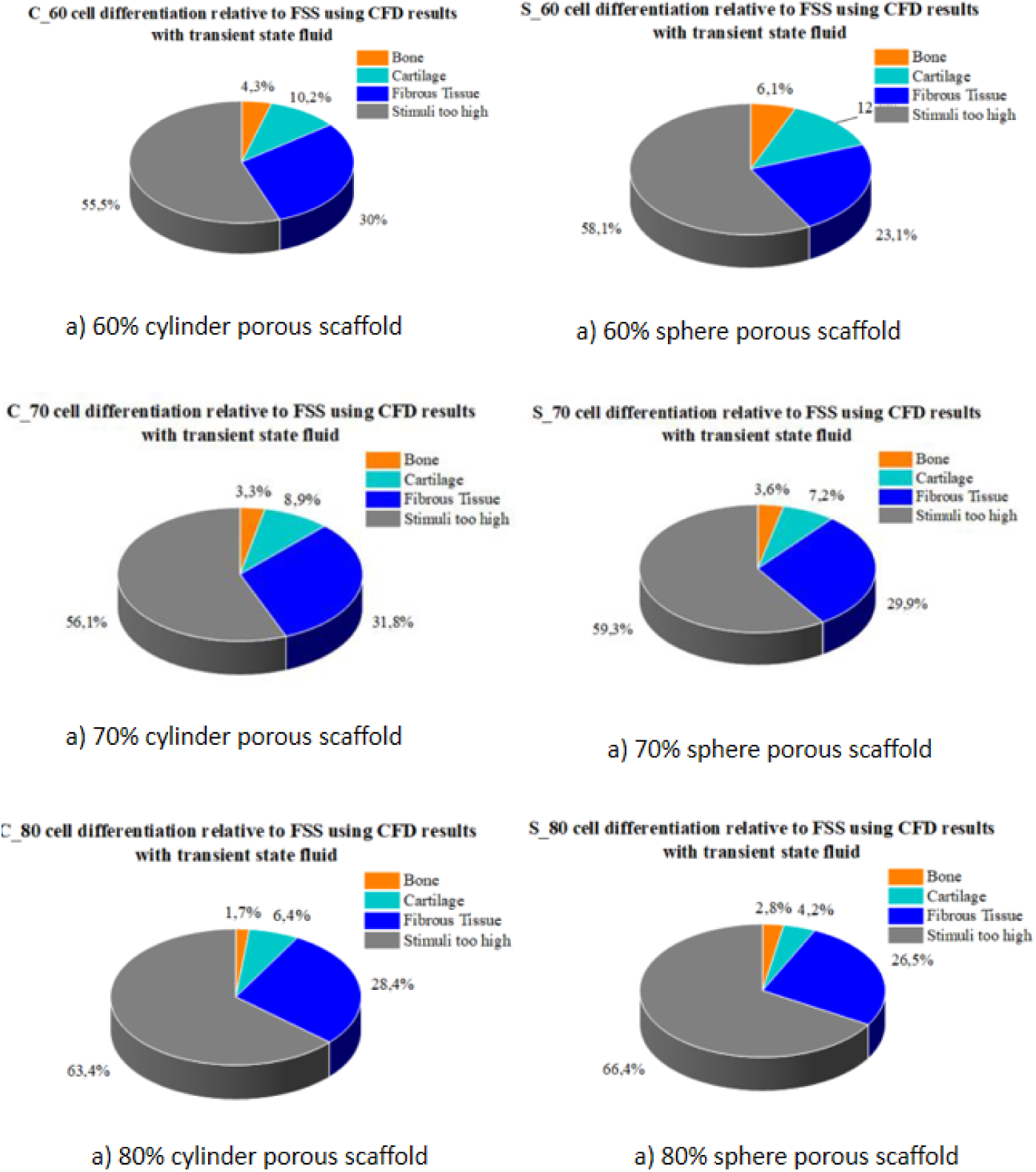
Cell differentiation analysis relative to Fluid Shear Stress (FSS) results in the superficial nodes using Computational Fluid Dynamics (CFD) at steady state inlet fluid, being bone differentiation orange, cartilage differentiation light blue, fibrous tissue differentiation dark blue, and stimuli too high to promote tissue differentiation.

**A.6.**
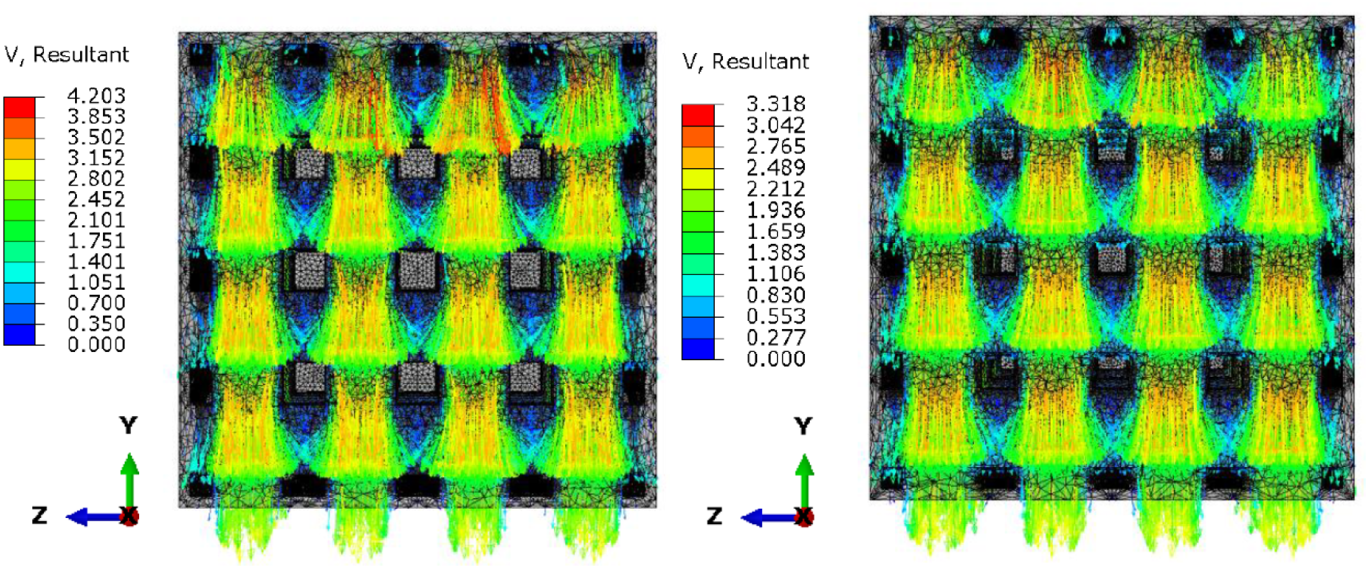
Computational Fluid Dynamics (CFD) simulation results showing the velocity vector map results at maximum fluid perfusion equal to 1 mm/s.

**A.7.**
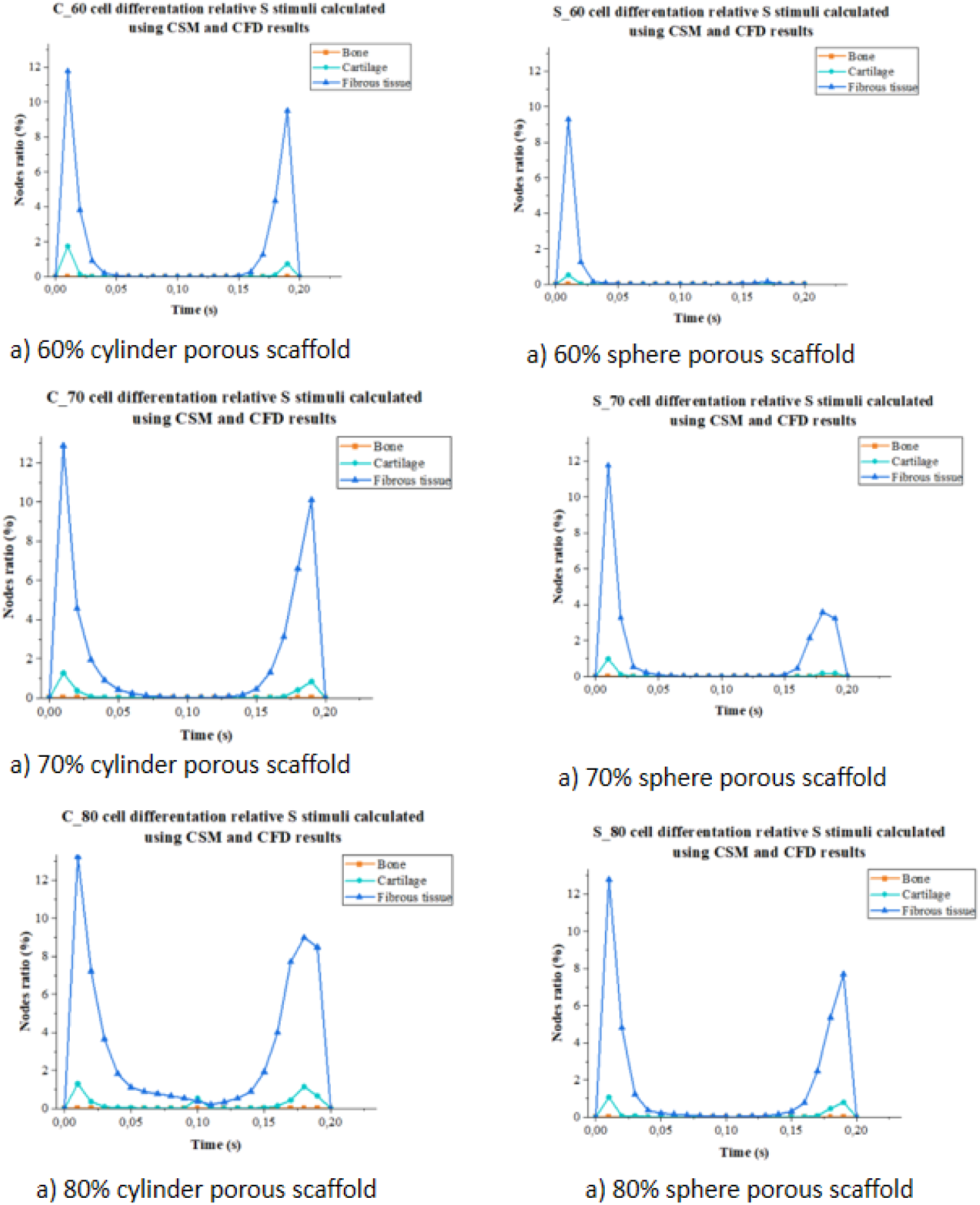
Cell differentiation analysis relative to S stimuli calculated with Equation 3, using the Fluid Shear Stress (FSS) results obtained with Computational Fluid Dynamics (CFD) models at transient state fluid perfusion and Shear Strain (SS) results obtained with Computational Solid Mechanics (CSM) models at dynamic compression. Rescaled axis and presenting bone differentiation orange, cartilage differentiation light blue, and fibrous tissue differentiation dark blue.

**A.8.**
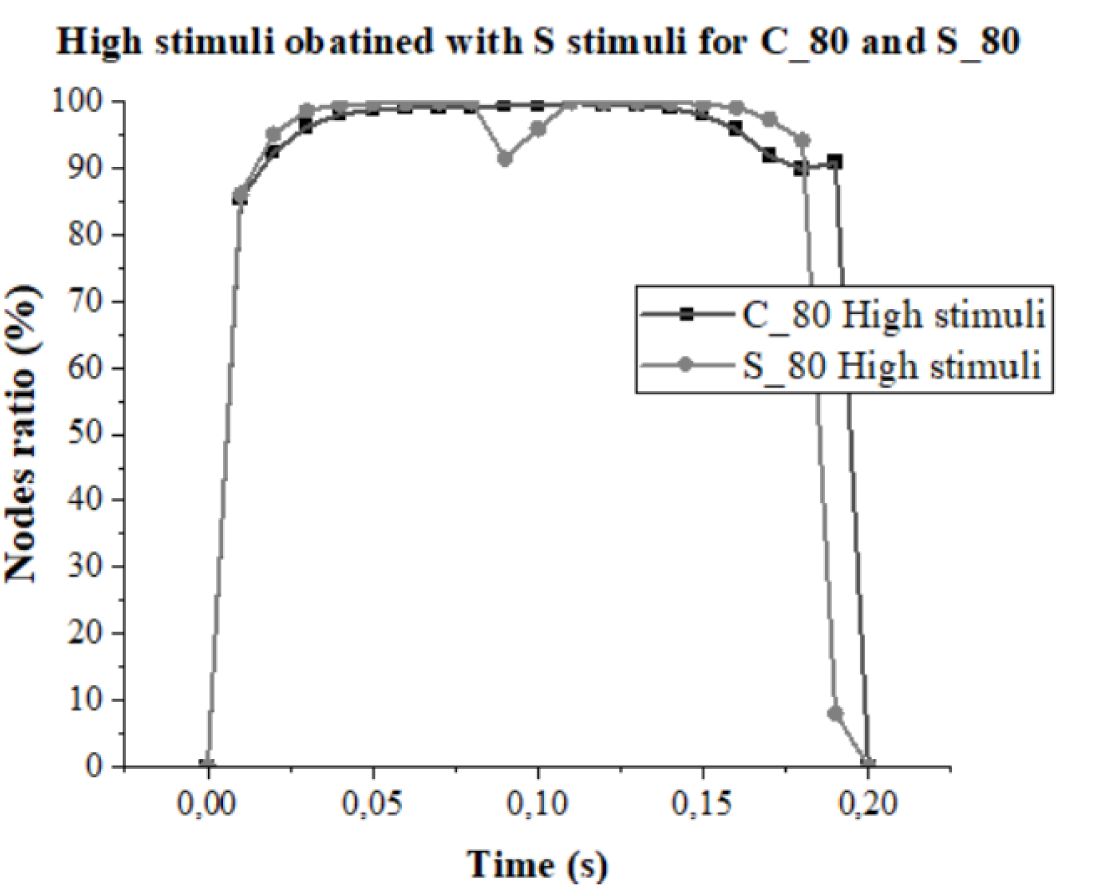
High stimuli obtained after calculating S stimuli using Fluid Shear Stress (FSS) obtained performing Computational Fluid Dynamics (CFD) and Shear Strain (SS) obtained performing Computational Solid Mechanics (CSM) simulation. 80% cylinder porous scaffold is presented in black and 80% sphere porous scaffold in grey.

**A.9.**
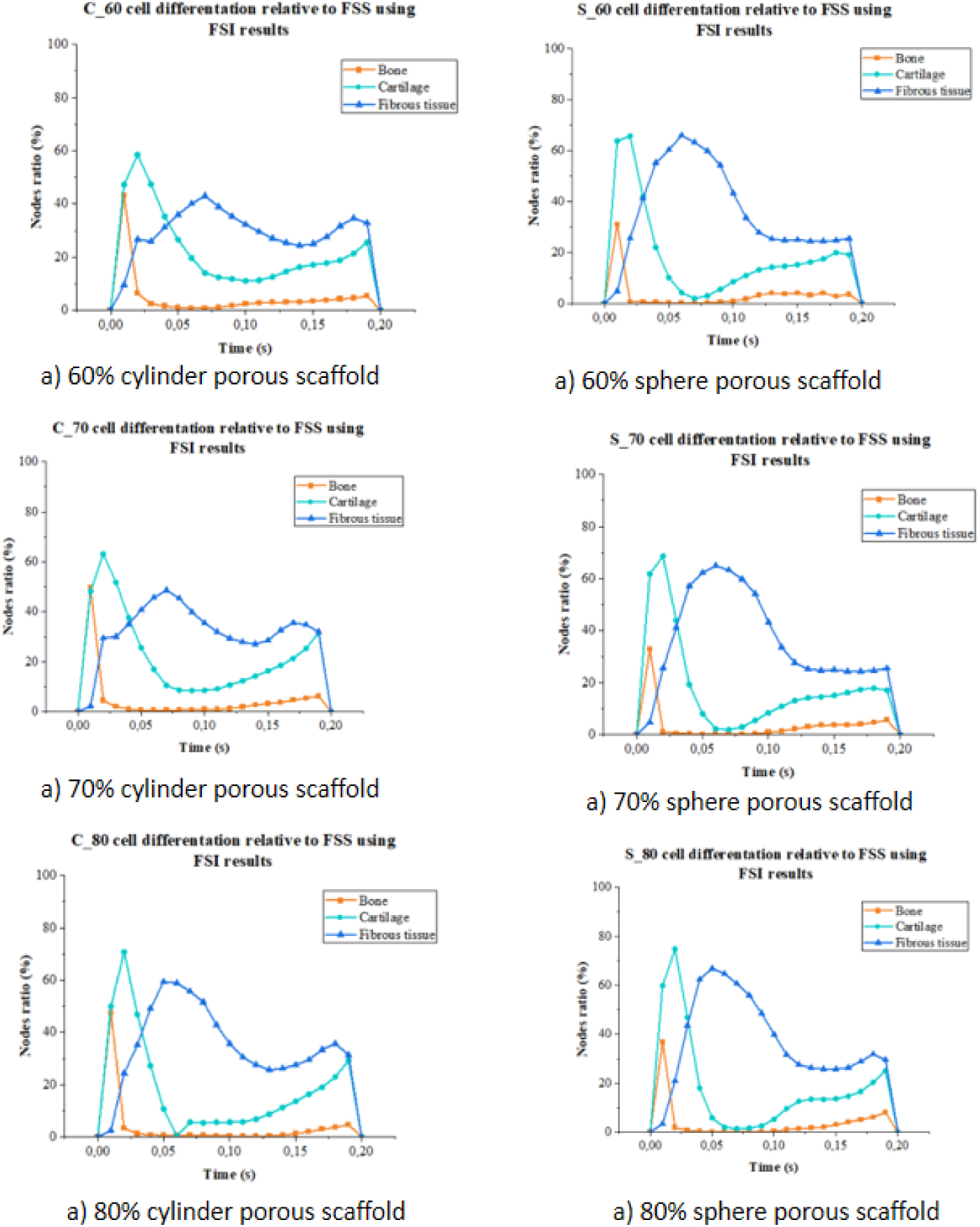
Cell differentiation analysis relative to Fluid Shear Strain (FSS) results in the superficial nodes using Fluid-Structure Interaction (FSI) at dynamic transient state fluid profile with no compression, being bone differentiation orange, cartilage differentiation light blue and fibrous tissue differentiation dark blue.

**A.10.**
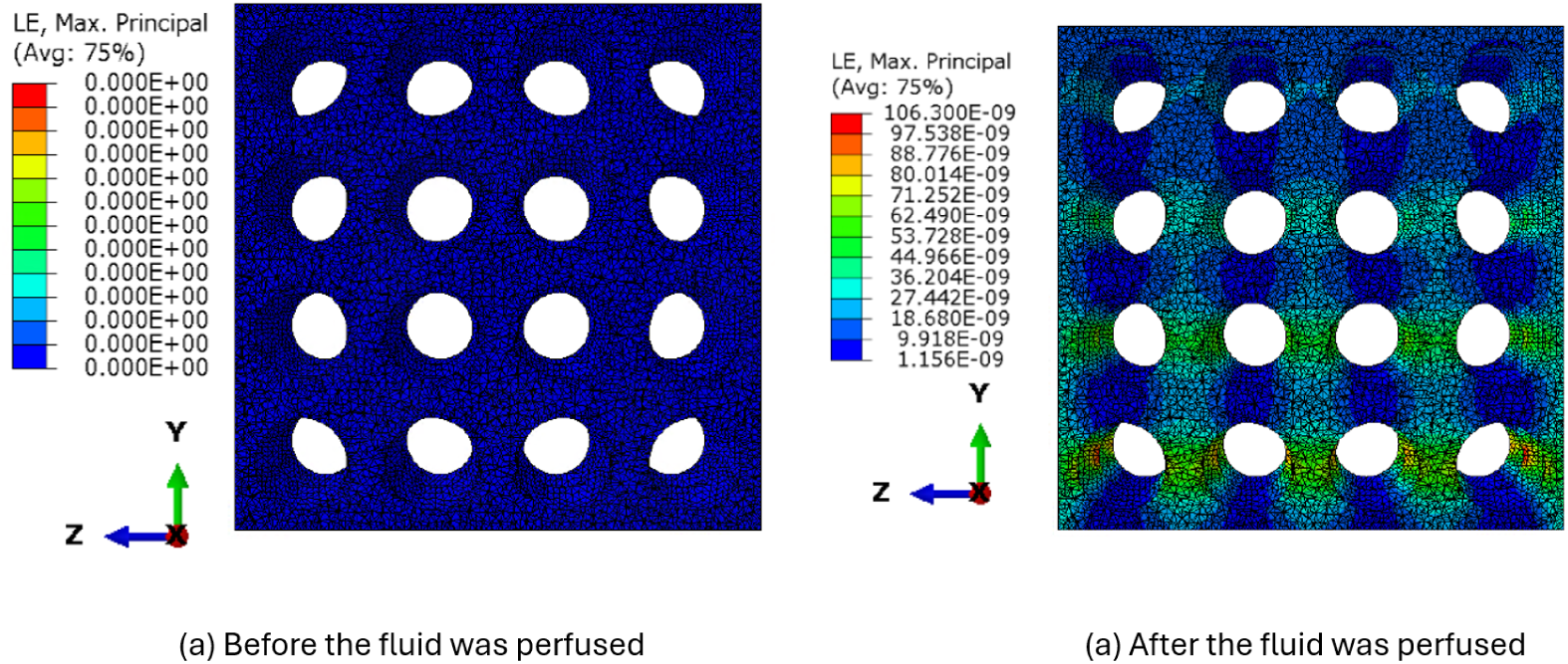
Shear strain (SS) color map of 80% sphere porous scaffold caused by the fluid stimuli.

**A.11.**
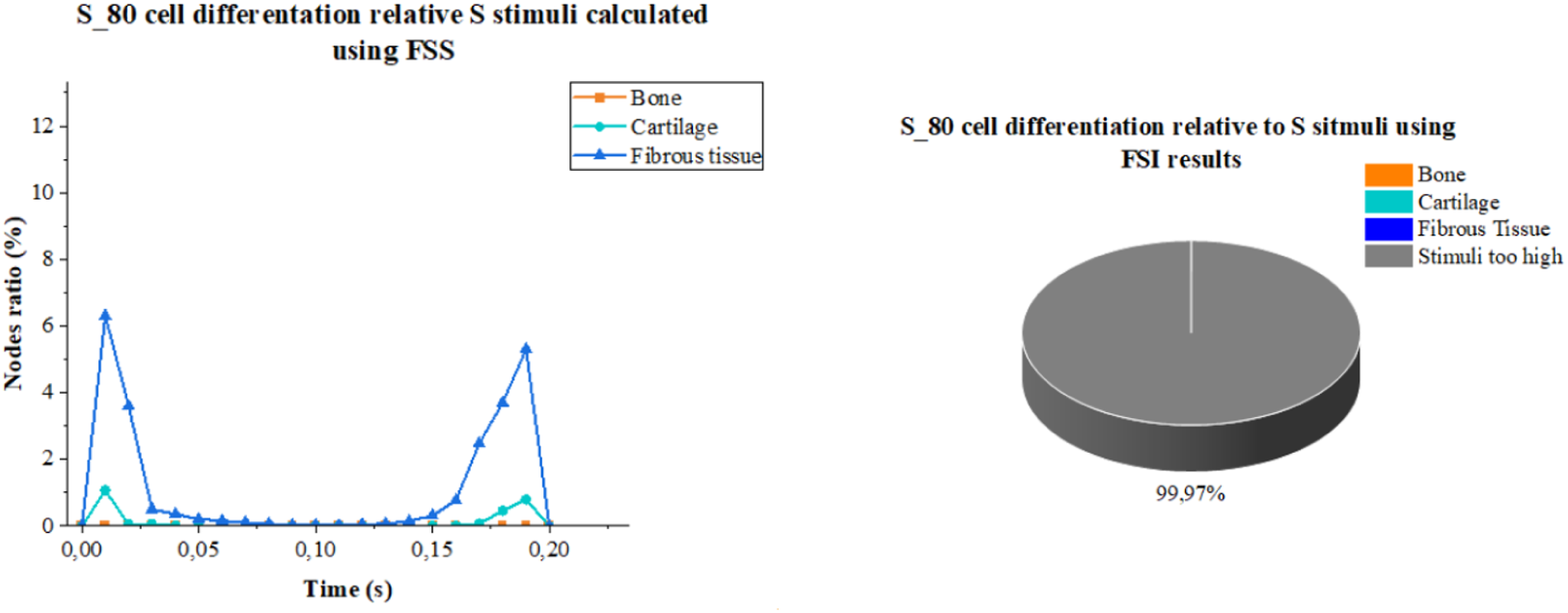
Cell differentiation results relative to S stimuli obtained implementing a Fluid-Structure Interaction (FSI) model with 80% sphere porous scaffold.

**A.12.**
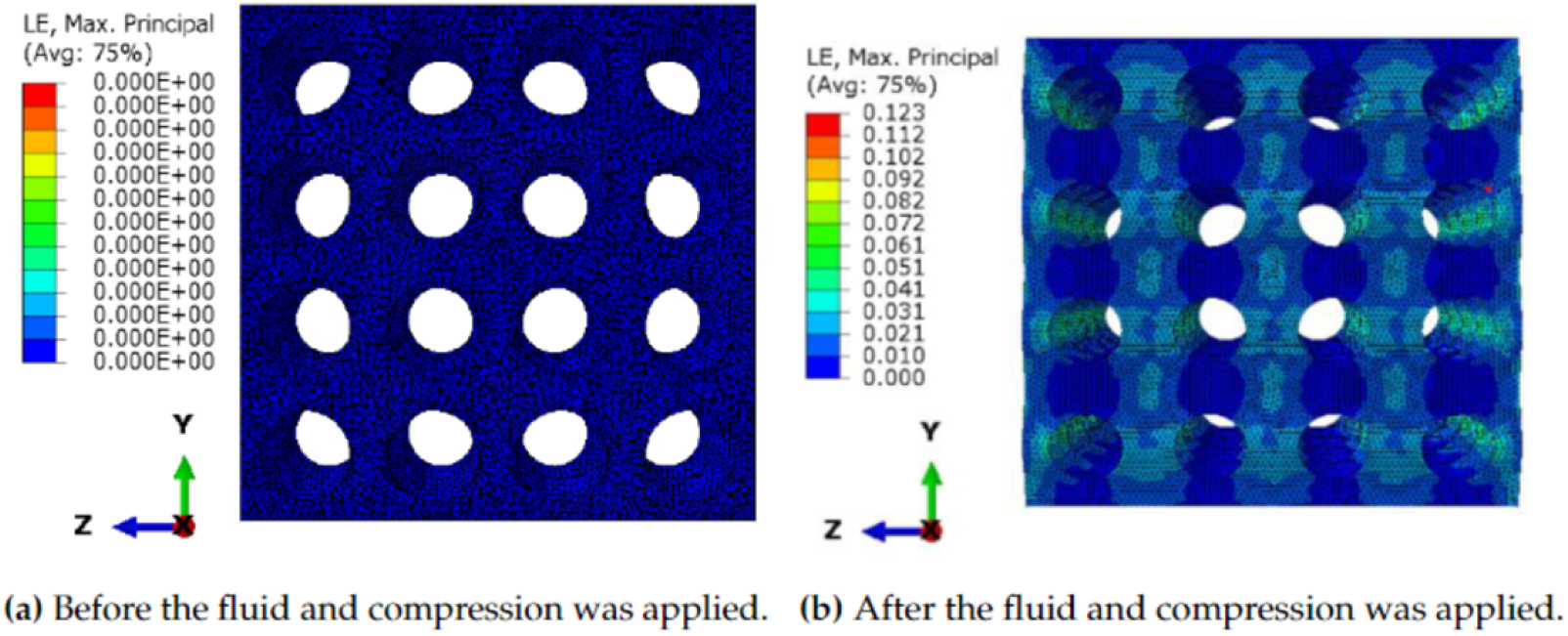
Shear Strain (SS) color map of 80% sphere porous scaffold caused by steady-state fluid perfusion at 1 mm/s and static compression of 5%.

## References

1. T. Adachi, Y. Osako, M. Tanaka, M. Hojo, and S. Hollister. Framework for optimal design of porous scaffold microstructure by computational simulation of bone regeneration. Biomaterials, 27:3964–3972, 2006.

2. T. Adachi, Y. Osako, M. Tanaka, M. Hojo, and S. J. Hollister. Framework for optimal design of porous scaffold microstructure by computational simulation of bone regeneration. Biomaterials, 27(21):3964–72, jul 2006.

3. D. Ali and S. Sen. Computational fluid dynamics study of the effects of surface roughness on permeability and fluid flow-induced wall shear stress in scaffolds. Annals Of Biomedical Engineering, 46:2023–2035, 2018.

4. A. Amini, C. Laurencin, and S. Nukavarapu. Bone tissue engineering: recent advances and challenges. Crit Rev Biomed Eng, 40:363–408, 2012.

5. P. Azizi, C. Drobek, S. Budday, and H. Seitz. Simulating the mechanical stimulation of cells on a porous hydrogel scaffold using an fsi model to predict cell differentiation. Frontiers in Bioengineering and Biotechnology, 11, 9 2023.

6. T. Casalini, F. Rossi, A. Castrovinci, and G. Perale. A perspective on polylactic acid-based polymers use for nanoparticles synthesis and applications. Frontiers In Bioengineering And Biotechnology, 7, 2019.

7. N. Castro, S. Ribeiro, M. M. Fernandes, C. Ribeiro, V. Cardoso, V. Correia, R. Minguez, and S. Lanceros-Mendez. Physically active bioreactors for tissue engineering applications, 10 2020.

8. C. W. F.J. Trueba. Changes in cell diameter during the division cycle of escherichia coli. Journal of Bacteriology, 142, 1980.

9. M. Fu, F. Wang, and G. Lin. Design and research of bone repair scaffold based on two-way fluid-structure interaction. Computer Methods And Programs In Biomedicine, 204:106055, 2021.

10. C. Geuzaine and J.-F. Remacle. Gmsh: A 3-d finite element mesh generator with built-in pre- and post-processing facilities. Int. J. Numer. Methods Eng, 79, 2009.

11. S. Hollister. Porous scaffold design for tissue engineering. Nature Materials, 4:518–524, 2005.

12. D. Lacroix and P. Prendergast. A mechano-regulation model for tissue differentiation during fracture healing: analysis of gap size and loading. Journal Of Biomechanics, 35:1163–1171, 2002.

13. D. Lacroix and P. J. Prendergast. A mechano-regulation model for tissue differentiation during fracture healing: analysis of gap size and loading. Journal of biomechanics, 35(9):1163–71, sep 2002.

14. G. L. M. Fu, F. Wang. Design and research of bone repair scaffold based on two-way fluid-structure interaction. Computer Methods and Programs in Biomedicine, 204, 2021.

15. M. Mohammadi, E. Rezabeigi, J. Bertram, B. Marelli, R. Gendron, S. Nazhat, and M. Bureau. Poly(d,l-lactic acid) composite foams containing phosphate glass particles produced via solid-state foaming using co2 for bone tissue engineering applications. Polymers, 12:231, 2001.

16. F. Morgan, L. Barnes, and T. Einhorn. Chapter 1 - the bone organ system: Form and function. pages 3–20. Fourth edition edition, 2013.

17. A. Olivares, E. Marsal, J. Planell, and D. Lacroix. Finite element study of scaffold architecture design and culture conditions for tissue engineering. Biomaterials, 30:6142–6149., 2009.

18. A. L. Olivares and D. Lacroix. Computational Methods in the Modeling of Scaffolds for Tissue Engineering. Springer-Verlag Berlin Heidelberg 2012, 2012.

19. P. Ouyang, H. Dong, X. He, X. Cai, Y. Wang, J. Li, H. Li, and Z. Jin. Hydromechanical mechanism behind the effect of pore size of porous titanium scaffolds on osteoblast response and bone ingrowth. Materials & Design, 183:108151, 2019.

20. M. Persson, P. Lehenkari, L. Berglin, S. Turunen, M. Finnilä, J. Risteli, M. Skrifvars, and J. Tuukkanen. Osteogenic differentiation of human mesenchymal stem cells in a 3d woven scaffold. Scientific reports, 8:10457, 2006.

21. C. Sandino and D. Lacroix. A dynamical study of the mechanical stimuli and tissue differentiation within a CaP scaffold based on micro-CT finite element models. 2010.

22. E. Schemitsch. Size matters: Defining critical in bone defect size! Journal Of Orthopaedic Trauma, 31(*5*), pages 20–22, 2017.

23. F. Sun, X. Sun, H. Wang, C. Li, Y. Zhao, J. Tian, and Y. Lin. Application of 3d-printed, plga-based scaffolds in bone tissue engineering. International journal of molecular sciences, 23:5831, 2022.

24. Y. Sun, B. Wan, R. Wang, B. Zhang, P. Luo, and D. Wang. Mechanical stimulation on mesenchymal stem cells and surrounding microenvironments in bone regeneration: Regulations and applications. Frontiers In Cell And Developmental Biology, 10, 2022.

25. X. Wang, L. Zhao, J. Fuh, and H. Lee. Effect of porosity on mechanical properties of 3d printed polymers: Experiments and micromechanical modeling based on x-ray computed tomography analysis. Polymers, 11:1154, 2019.

26. J. Zeltinger, J. Sherwood, R. M. D. Graham, and L. Griffith. Effect of pore size and void fraction on cellular adhesion, proliferation, and matrix deposition. Tissue Engineering, 7:5, 2001.

27. F. Zhao, T. T. Vaughan, and L. McNamara. Quantification of fluid shear stress in bone tissue engineering scaffolds with spherical and cubical pore architectures. Biomechanics And Modeling In Mechanobiology, 15:561–577, 2015.

